# Competing gene regulatory networks drive naive and memory B cell differentiation

**DOI:** 10.1101/2025.06.04.657836

**Authors:** Pietro Demela, Laura Esposito, Pietro Marchesan, Davide Bolognini, Edoardo Giacopuzzi, Eugenia Ricciardelli, Paolo Ferrari, Silvia Bombelli, Clelia Peano, Daniele Prati, Luca Valenti, Blagoje Soskic

**Author notes:** Corresponding author: Blagoje Soskic.

## Abstract

Elucidating the gene regulatory networks (GRNs) that govern human B cell differentiation is essential for understanding immune responses to infection, vaccination and autoantigens. Here, we show that individual naive B cells can give rise to both plasma cells and germinal centre (GC) B cells. In contrast, memory B cells display a progressive increase in IRF4 activity over time, leading to PRDM1 induction and exclusive differentiation into plasma cells. Using CRISPR-based perturbations, we demonstrate that IRF4 is indispensable for both GC and plasma cell development. Notably, IRF4 promotes GC fate independently of PRDM1, as PRDM1 disruption did not impair GC differentiation. We also find that while the abundance of antibody mRNAs is clonally correlated, class switch recombination (CSR) occurs stochastically and is clonally independent. Together, these findings reveal distinct regulatory dynamics during naive and memory B cell activation and offer new insights into the GRNs underlying human B cell fate decisions.

## Introduction

Upon encountering an antigen, naive B cells proliferate and their progeny can differentiate into germinal centre (GC) cells, memory B cells, short-lived plasmablasts, and long-lived plasma cells, each playing a distinct role in defending the organism against pathogens (Inoue and Kurosaki 2024; Nutt et al. 2015; De Silva and Klein 2015; Victora and Nussenzweig 2022; Cyster and Allen 2019). Within the GC, B cells diversify their antibodies to respond to a wide array of pathogens. To enhance the diversity of the antibody repertoire, B cells undergo two key processes: somatic hypermutation (SHM) and class switch recombination (CSR) (De Silva and Klein 2015; Z. Xu et al. 2012; Allen, Okada, and Cyster 2007; Victora and Nussenzweig 2022). SHM improves antigen binding via point mutations in V(D)J segments of the immunoglobulin gene, while CSR replaces the constant region to alter antibody isotype. CSR generates antibodies with different effector functions (IgG, IgA or IgE) while preserving their original antigen specificity. Even though the molecular machinery responsible for CSR has been largely known (Z. Xu et al. 2012), the causes of differential CSR outcomes are still unclear. Specifically, in a work by Horton and colleagues, CSR outcomes were explained assuming an underlying stochastic mechanism (Horton et al. 2022). In another study, intrinsic transcriptional heterogeneity of naive B cells was indicated as the leading cause of differential isotype switching outcomes (Y. L. Wu et al. 2017). The dynamics and probability of antibody class switching have been largely studied in mouse models by isolating switched and unswitched cells and comparing their proportions and phenotype before and after antigen exposure. Technical advances in single-cell sequencing that allow exploration of both gene expression and antibody repertoire provide a powerful approach to capturing the transcriptomic landscapes of individual human B cells, their progeny and their antibody repertoire, allowing us to investigate transcriptional dynamics involved in this critical decision point of the immune response.

While naive B cells are critical in the primary response to antigen, memory B cells are particularly important during a secondary infection. Upon activation, memory B cells rapidly give rise to plasmablasts (Arpin, Banchereau, and Liu 1997), that increase circulating levels of high-affinity antibodies, leading to more efficient clearance of the pathogen (Inoue and Kurosaki 2024; Lam, Lee, and Farber 2024). The heightened responses of memory B cells could be partially attributed to the antibodies they express, which have already undergone diversification via SHM and/or CSR and exhibit high affinity for the antigen (Inoue and Kurosaki 2024; Mesin et al. 2020; Gitlin et al. 2016). This increased affinity may lower the activation threshold of memory B cells (Ambegaonkar et al. 2024; Anderson et al. 2009; Shao et al. 2024). However, experiments with naive B cells engineered to express memory-derived antibodies, showed that these cells responded similarly to wild-type naive B cells (Kometani et al. 2013). Thus, the heightened responses of memory B cells cannot be explained only by their higher antigen affinity. Finally, clonal analysis of memory B cells in mice, showed that secondary GCs are composed predominantly of clones without primary GC experience, and thus likely naive in origin (Mesin et al. 2020). As such, re-enter and new diversification of antibodies in previously diversified memory B cells is a rare event (Mesin et al. 2020; Li et al. 2024; Schiepers et al. 2024). It has been speculated that the dominance of naive-derived cells in secondary GCs may be due to precursor abundance rather than suppression of MBCs - i.e. a number of different naive B cells with high enough affinity to initiate a primary GC reaction is larger than the number of memory cells (Mesin et al. 2020; Tas et al. 2022). Another possibility is that high-affinity antibodies limit GC entry for targeted clones (Schiepers et al. 2024; Tas et al. 2022). Importantly, these studies have not investigated whether differentiation differences can also be attributed to the cell-intrinsic phenomenon of human naive and memory B cells, and what are differentiation trajectories in the absence of competition or selection.

B cell activation is mainly controlled by two opposite genetic circuits (Nutt et al. 2015; Sciammas et al. 2011; H. Xu et al. 2015). High levels of IRF4 induce PRDM1 expression to dictate the differentiation of B cells towards the plasma cell fate (Ochiai et al. 2013). On the contrary, when the levels of IRF4 are low, BCL6 and BACH2 steer B cell responses towards the GC fate, where B cells proliferate and diversify their antibodies. PRDM1 and BCL6 inhibit each other and the balance between them dictates which cell state B cell will acquire (Shao et al. 2024; Nutt et al. 2015; Sciammas et al. 2011; Ochiai et al. 2024, 2013; Kometani et al. 2013; H. Xu et al. 2015; Scharer et al. 2020). However, it is still unclear how naive and memory B cells differently activate these opposite gene programs networks, and which downstream genes are in turn regulated. An in-depth understanding of how human naive and memory B cell activation is regulated over time is critical in understanding pathogen responses and in turn increase the effectiveness of vaccination and therapies.

To investigate the gene programs involved in B cell activation and antibody switching, we profiled the gene expression at the single cell level as well as the antibody repertoire of human naive and memory B cells at resting and following multiple time points of activation. We showed that individual naive B cells can give rise to both plasma cells and GC cells, with expression of key differentiation genes tightly regulated in a clonal manner. Despite clonal regulation of antibody transcript levels, CSR occurs independently and stochastically. We also showed that memory B cells exhibit progressive IRF4 activation, culminating in PRDM1 expression and plasma cell differentiation. Finally, through CRISPR perturbations, we defined IRF4 as a central regulator of both GC and plasma cell fates, acting independently of PRDM1 in GC commitment. These findings highlight the distinct and dynamic regulatory circuits guiding naive and memory B cell differentiation outcomes and advance our understanding of the GRNs orchestrating human B cell activation.

## Results

### Naive and memory B cells have distinct differentiation outcomes

To investigate the activation dynamics, differentiation outcomes and control of CSR in naïve and memory B cells, we established an *in vitro* model of B cell activation. We purified human naive and memory B cells from PBMCs (Methods). To elucidate GRNs driving isotype switching, we removed B cells expressing switched antibody isotypes. Therefore, here naive B cells are defined as CD20^+^ CD27^-^ IgG^-^ IgA^-^ and unswitched memory B cells are defined as CD20^+^ CD27^+^ IgG^-^ IgA^-^ (**Supplementary Fig. 1a**). Next, we optimised a stimulation cocktail that models the T-dependent B cell activation (**Supplementary Fig. 1b-c**). We used an IgM cross-linker providing BCR stimulation, multimeric CD40L that engages costimulatory receptor CD40 on the B cell, and cytokines IL-2 and IL-21 (Methods).

To comprehensively characterise activation dynamics of naive and memory B cells, we performed single-cell RNA sequencing (scRNA-seq) and B cell receptor sequencing (BCR-seq) in resting cells (day 0), before the first division (day 1), at the time of the first division (day 3) and following the full B cell differentiation (day 6) (**Figure 1a**). Following quality control (Methods), we retained 118,891 cells. We used uniform manifold approximation and projection (UMAP) for dimensionality reduction and observed that cells separated by activation time point and cell type (**Figure 1b**). This indicated that the transcriptional response to activation is both dynamic and cell type-specific. To further elucidate the molecular differences between naive and memory B cells, we performed differential gene expression analysis. This revealed distinct transcriptional responses following activation, with 1537, 1275, 1159 and 3017 genes significantly differentially expressed between the two cell types across days 0, 1, 3 and 6, respectively (**Figure 1c, Supplementary Table 1**). On day 1, both naive and memory cells upregulated costimulatory receptor *CD40* (**Figure 1d**), indicating early activation and preparation for efficient interaction with T cells expressing CD40L. On the other hand, a proinflammatory chemokine *CCL22* is highly expressed in memory B cells on day 1 and day 3 but not in naive cells (**Figure 1d**). Likewise, *CCL17* showed specific expression in day 3 memory B cells (**Figure 1d**). CCL17 and CCL22 have been shown as critical in recruiting CCR4-expressing T cells, suggesting that memory B cells are more efficient in recruiting T cells (Liu et al. 2021). In contrast, *ID3,* which has been demonstrated to promote the differentiation of GC B cells and negatively regulate plasma cell differentiation (Gloury et al. 2016; Chen et al. 2016), showed markedly higher expression in activated naive cells (**Figure 1d**).

**Figure 1.**
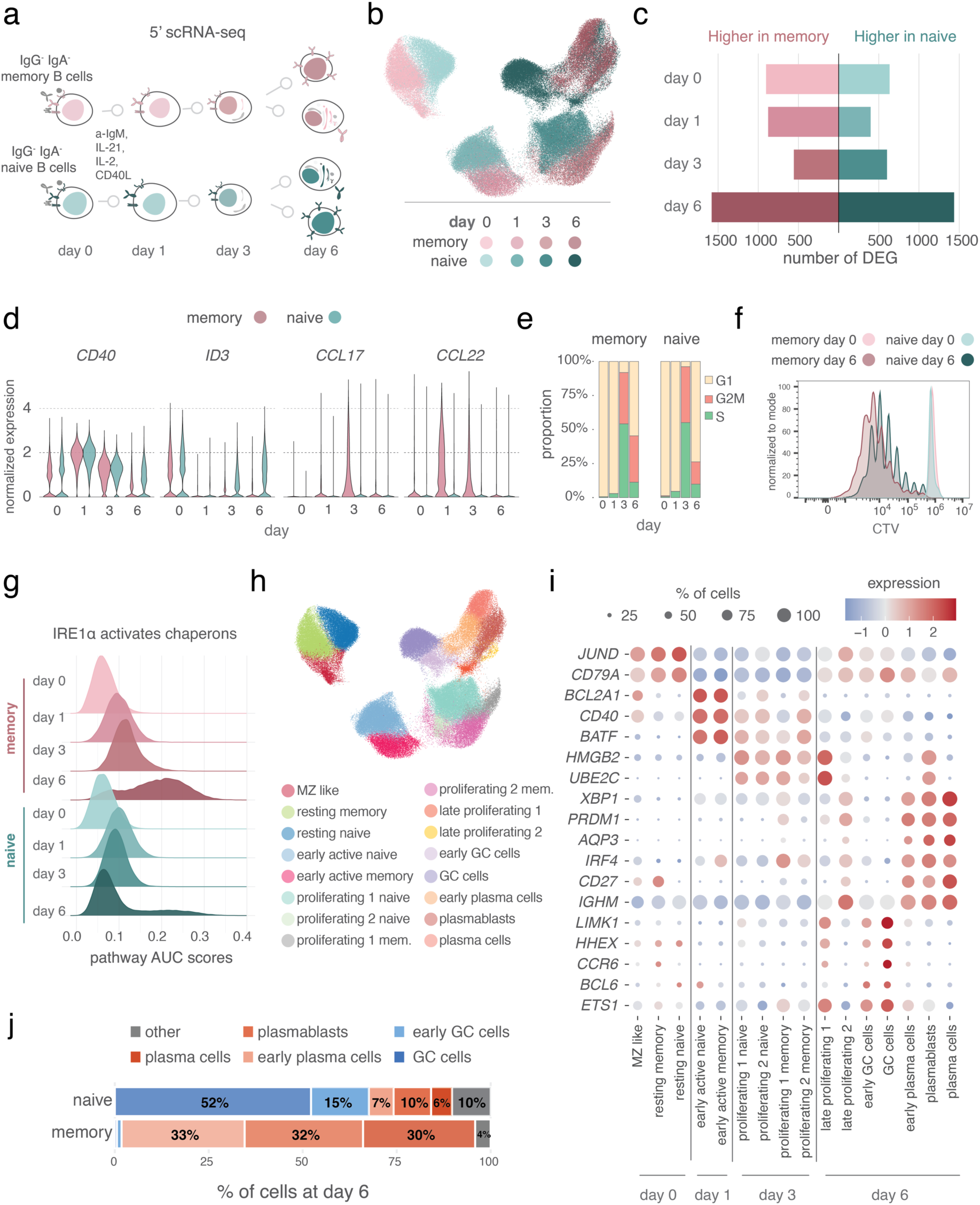
Naive and memory B cells have distinct differentiation outcomes. **a**) Schematic overview of the study design. **b**) UMAP embeddings of scRNA-seq of naive and memory B cells at resting and 1, 3 and 6 days following stimulation. Colours represent cell type and time point. **c**) Barplot illustrating the number of differentially expressed genes (DEG) between naive and memory B cells at each time point. **d**) Normalized expression level of *CD40, ID3, CCL22* and *CCL17*. **e**) Proportion of cells in different cell cycle phases by cell type and time point. **f**) CellTrace violet (CTV) staining of naive and memory B cells at day six. **g**) Density plot of AUCell scores for the “IRE1α activates chaperons” pathway stratified by cell type and time point. **h**) UMAP embeddings of scRNA-seq of naive and memory B cells at resting and 1, 3 and 6 days following stimulation, coloured by cell states. **i**) Dot plot of cell state specific genes. Scaled gene expression is represented on a blue-to-red gradient: blue indicates below-average expression, red indicates above-average expression. Dot size reflects the proportion of cells expressing each gene. **j**) Proportion of cells in different cell states for naive and memory cells at day 6, colours represent cell states. a,b,c,d,f,g) shades of purple and green represent memory and naive B cells, respectively.

Both cell types showed a progressive increase in cell cycle activity, peaking at day 3, when approximately 90% of cells were in S or G2M phases in both cell types (**Figure 1e)**. By day 6, however, approximately 70% of naive and 50% of memory cells had returned to the G1 phase, indicating a reduction in proliferation and increased differentiation. This observation is consistent with the higher proportion of memory B cells in later divisions (**Figure 1f**) and overall higher number of memory B cells than naive B cells at day 6 (**Supplementary Fig.1c**).

We noted the highest number of differentially expressed genes between naive and memory B cells at day 6 (**Figure 1c, Supplementary Table 1**). Upregulated genes in memory B were significantly enriched for pathways and signatures related to endoplasmic reticulum (ER) stress response (**Figure 1g, Supplementary Table 2**). This included pathways such as “IRE1α activates chaperones”, “XBP1(S) activates chaperone genes“,”Unfolded Protein Response (UPR)” and “Transport to the Golgi and subsequent modification” (**Figure 1g**). As these signatures are characteristic of plasma cell differentiation (Nutt et al. 2015), we hypothesised that memory B cells are more efficient in differentiating into plasma cells than naive B cells. Therefore, to further dissect the heterogeneity of these populations, we performed unsupervised clustering of B cells and identified 16 cell states (**Figure 1h-i**). Strikingly, memory B cells almost exclusively differentiated into plasmablasts and plasma cells (**Figure 1i-j**). Unlike plasma cells, plasmablasts were predominantly in S or G2M phases of the cell cycle (**Supplementary Fig.1d**), and expressed a lower amount of *IGHM* antibody than plasma cells (**Supplementary Fig. 1e**). In contrast, naive B cells had a bifurcated differentiation and generated two distinct cell states (**Figure 1j**). One differentiation branch resulted in plasmablasts and plasma cells that clustered with memory counterparts, while the other resulted in a germinal centre (GC) phenotype. This GC cluster was defined based on the high expression of GC markers, including *BCl6*, *CD81*, *LIMK1* and high expression of multiple activation-related genes, and the absence of plasma cell markers (e.g. *XBP1, IRF4, PRDM1* and low *IGHM*) (**Figure 1i**). These cells also expressed markers of memory precursor state, such as *CCR6* and *HHEX* (Laidlaw et al. 2020; Suan et al. 2017) suggesting that they contain the source of new memory B cell population. Differential gene expression analysis between plasma cells and GCs identified more than 8,000 differentially expressed genes, suggesting a significant fate divergence. (**Supplementary Table 3**).

Taken together, our data demonstrate that following the same stimulation, naive and memory B cells upregulate distinct genes and pathways, resulting in significant divergence in the end cell states.

### Opposing transcription factor activities regulate fate decision

Next, we hypothesised that the divergent differentiation outcomes of naive and memory B cells are driven by differences in transcription factor activity across stimulation. We reasoned that a cell’s identity is a consequence of GRNs in which a defined set of TFs regulate downstream targets. We investigated changes in activity of transcription factors and their target genes which are referred to as regulons. In order to infer GRN activity during differentiation of naive and memory B cells we applied SCENIC (Aibar et al. 2017). Briefly we identified modules of co-expressed genes and filtered them based on the presence of transcription factor-binding motifs in their promoters. At the resting state (day 0), naive and memory B cells had highly similar regulon activity for the vast majority of TFs (r = 0.89, p-value = 3.2×10^-37^) (**Figure 2a, Supplementary Fig. 2a, Supplementary Table 4**). Among a few differences we observed FOXO1 and FOXO3, both TFs were previously implicated in B cell development and BCR signalling (Ottens et al. 2018; Dengler et al. 2008). Overall, the high degree of similarity of GRNs in the resting state is not surprising given that naive and memory B cells belong to the same cell type and share many core biological functions. However, TF activity became increasingly distinct between naive and memory B cells as the activation progressed, culminating on day 6 (r=0.4) (**Figure 2a, Supplementary Fig. 2a and 2b**). A large number of regulons appeared to exhibit opposing activity in naive and memory cells on day 6. Notably, IRF4 regulon activity (i.e. high expression of IRF4 target genes) showed stronger activity in day 6 activated memory B cells compared to naive, while SPI1 (PU.1) regulon was most active in naive cells (**Figure 2b, Supplementary Fig. 2a**). These two regulons were anti-correlated and mapped onto distinct fate trajectories. While IRF4 regulon activity was strongly associated with the plasma cell fate, SPI1 activity characterized the GC state (**Figure 2b, Supplementary Fig. 2b**). Cells with high IRF4 activity also showed high XBP1 activity (**Figure 2b**) and high antibody expression (**Figure 2c**) confirming their identity as plasma cells.

**Figure 2.**
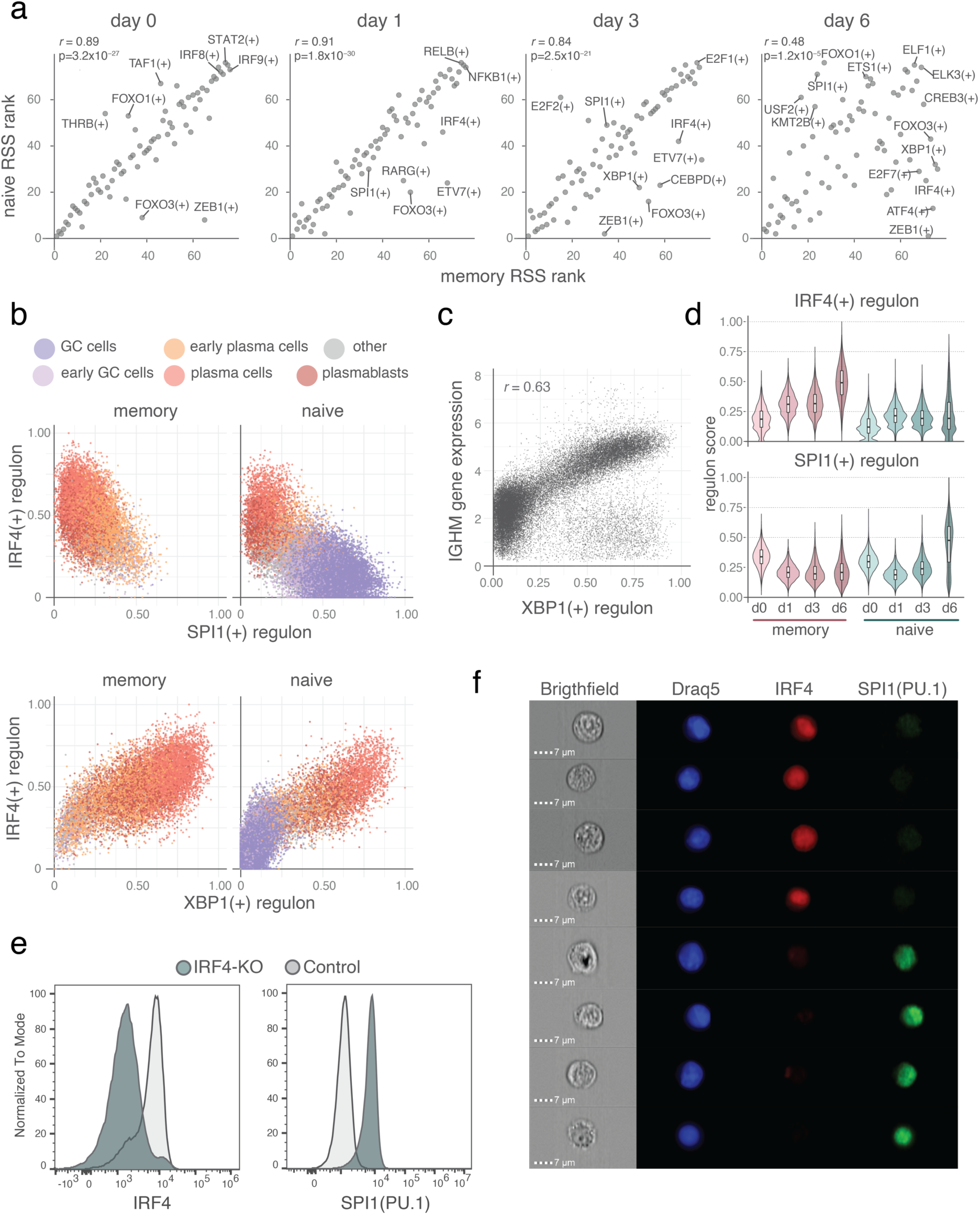
Opposing transcription factor activities regulate fate decision. **a**) Comparison of regulon specificity score (RSS) rankings between naive and memory B cells across time points. For each condition and cell type, RSS are ranked from lowest to highest. Pearson correlation coefficient (r) and its two-sided test P values are shown. **b**) Dot plots showing the SPI1, IRF4 and XBP1 regulon scores in day six naive and memory B cells. Colours represent cell states. **c**) Dotplot showing the correlation between XBP1 regulon scores and normalized *IGHM* gene expression in naive and memory B cells at day 6. **d**) IRF4 and SPI1 regulon scores across time points for naive and memory B cells. Shades of purple and green represent memory and naive B cells, respectively. **e**) Histogram showing intracellular staining of IRF4 and PU.1 (SPI1) in IRF4 knock-out activated naive B cells. **f**) Representative ImageStream images of activated naive B cells at day six stained for nucleus (Draq5), IRF4 and SPI1. Images were acquired at 60× magnification.

Interestingly, IRF4 activity was already increased in memory B cells during early activation and continued to rise throughout the activation trajectory, consistent with the observation that memory B cells almost exclusively differentiate into plasma cells (**Figure 2d**). In contrast, in naive B cells we observed a transient increase in IRF4 activity during early activation, which was subsequently downregulated and remained high only in the subset of cells that differentiated into plasma cells (**Figure 2d**). Strikingly, the dynamics of SPI1 (PU.1) activity showed an inverse pattern (**Figure 2d**). In memory B cells, SPI1 activity was strongly suppressed upon activation and remained low over time. However, in naive B cells, following an initial downregulation, SPI1 activity increased progressively, peaking by day 6 of activation. This opposing behavior of IRF4 and SPI1 across naive and memory cells led us to hypothesize that IRF4 activity may inhibit SPI1 activity. To test this relationship, we used CRISPR–Cas9 to knock-out IRF4 in activated naive B cells. IRF4 deletion resulted in an increase in SPI1 (PU.1) expression, supporting the hypothesis that IRF4 negatively regulates SPI1 (**Figure 2e**). Finally, we performed imaging flow cytometry to assess the subcellular localization of IRF4 and SPI1 (PU.1). This analysis revealed a clear and mutually exclusive pattern. B cells that have IRF4 localised in the nucleus were negative for SPI1 (PU.1), and vice versa (**Figure 2f**).

Taken together, these findings demonstrate that transcription factor regulatory networks diverge significantly between naive and memory cells during activation. By day 6, opposing GRNs are active, reflecting the underlying commitment to either plasma cell or GC fates.

### Plasma cells and GC cells can arise from the activation of the same progenitor cell

Given that naive B cells differentiate into plasma cells and GC cells, we next sought to identify whether this bifurcation arises from a population-level behavior (i.e. different cells commit to different fates), or whether an individual cell is capable of generating both fates (**Figure 3a**). We reasoned that if one cell gives rise to both cell states at day 6, then as that cell divides and forms a clone, we should observe both cell states within that clone. To address this, we developed an improved B cell stimulation protocol that allowed us to expand a smaller number of naive B cells in culture (2,000 B cells at day 0). On day 6, we performed single-cell RNA-and BCR-sequencing on the entire culture, preserving both clonal diversity and size. We sequenced 23,484 cells, and identified 265 clones with at least 10 cells (**Figure 3b**). Strikingly, this revealed that a large proportion of the clones contained both plasma cell and GC cell states while others produced only plasma or GC cells (**Figure 3c**). Interestingly, we also noted that clones enriched in plasma cell fraction tended to be larger (**Figure 3d**). To validate this observation, we stained for XBP1 and IRF4 as proxies for plasma cell fate, and showed that XBP1^+^IRF4^+^ cells proliferate more extensively than XBP^-^ IRF4^-^ (**Figure 3e, Supplementary Fig. 3a**).

**Figure 3.**
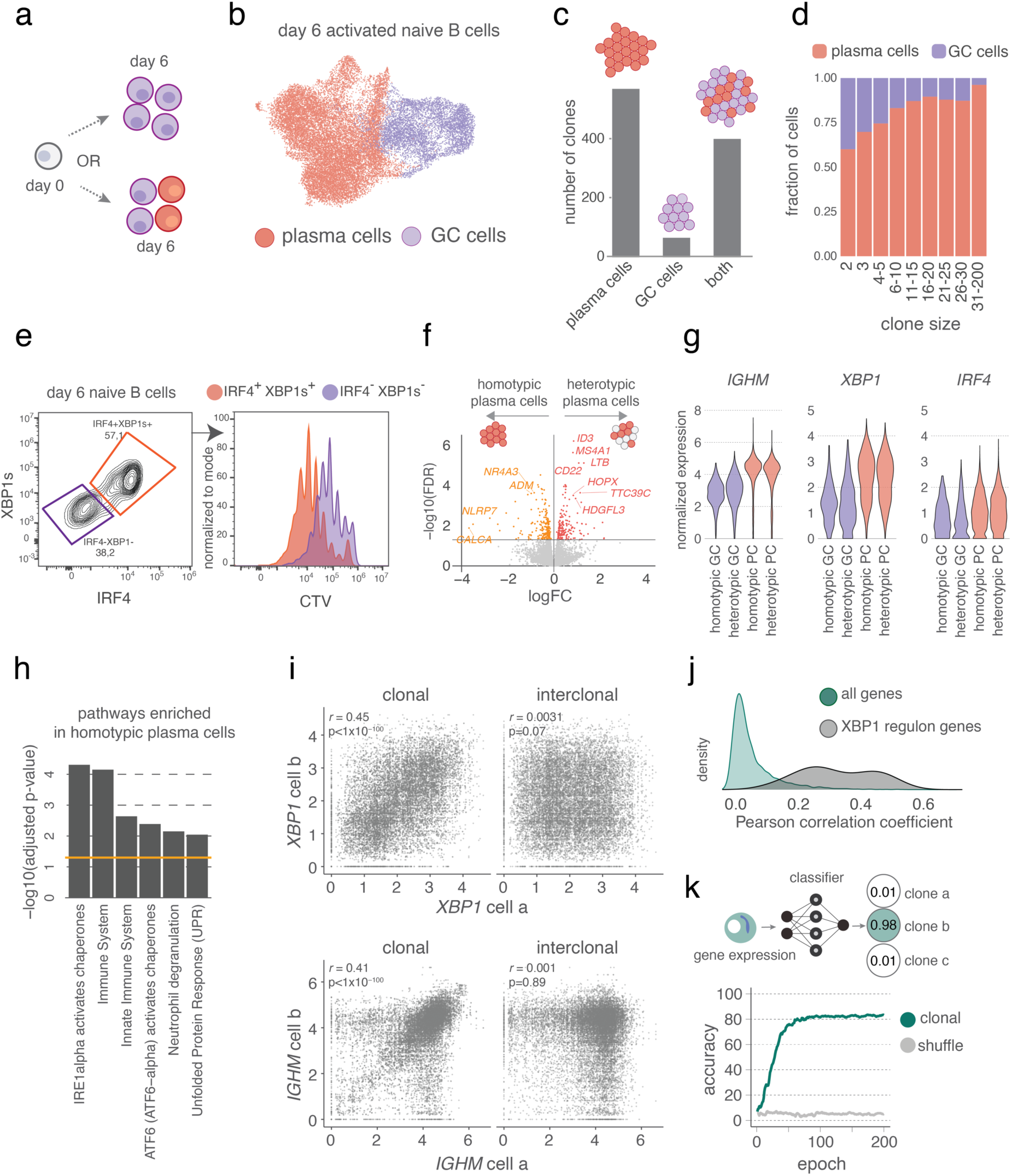
Plasma cells and GC cells can arise from the activation of the same progenitor cell. **a**) Schematic overview of two hypotheses. **b**) UMAP embeddings of scRNA-seq of naive B cells six days following stimulation. Colours represent cell state. **c**) Barplot of number of clones giving rise to only plasma cells, only GC cells or both. **d**) Proportion of different cell states per clonal size. Colours represent cell state. **e**) The left plot shows intracellular staining of IRF4 and XBP1 in naive B cells at day six after activation. The right plot shows CellTrace violet (CTV) proliferation analysis of the subpopulations on the left. **f**) Volcano plot of differential gene expression between homotypic and heterotypic plasma cells. Grey indicates non-differentially expressed genes; orange and red indicate genes up-and downregulated in heterotypic plasma cells, respectively. **g**) Normalized expression level of *IGHM, XBP1 and IRF4,* separated by clone type. Colors represent cell state. **h**) Top enriched reactome pathways in homotypic plasma cells. **i**) Interclonal and clonal correlation of *XBP1* and *IGHM* normalized gene expression. Pearson correlation coefficient (r) and its two-sided test P values are shown. **j**) Density plot of clonal Pearson correlation coefficient for XBP1 regulon genes compared to all other genes. **k**) Classification accuracy (%) of the classifier along training for the validation dataset. Accuracy is calculated as the proportion of correctly assigned cells to clones to total cells. a,b,c,d,g) orange and purple represent plasma cells and GC cells respectively.

We next assessed whether the plasma cells from “plasma cell only” clones were different from plasma cells in mixed cones (plasma cell+GC cells) (**Figure 3f)**. We referred to these populations as homotypic and heterotypic plasma cells respectively. Importantly, there was no difference in key plasma cell genes *IGHM, XBP1 and IRF4* further confirming their plasma cell identity (**Figure 3g**). However, 412 genes were differentially expressed between homotypic and heterotypic plasma cells. Genes upregulated in homotypic plasma cells were enriched in ER stress response pathways including “IRE1alpha activates chaperones”, “ATF6 (ATF6−alpha) activates chaperones” and “Unfolded Protein Response (UPR)” (**Figure 3h**). This demonstrated that plasma cells from homotypic clones have stronger ER stress signatures.

Next, we hypothesised that if heterotypic and homotypic plasma cells have distinct transcriptional profiles, each clone could carry its own unique gene expression signature. We reasoned that if the level of gene expression is heritable across cell divisions then sister cells should display similar mRNA expression profiles. To test this we analyzed the gene expression correlation between sister cells and compared them to correlations observed in random cell pairs (**Supplementary Fig. 3b**). Strikingly many top correlated genes were involved in plasma cell differentiation and function. For examples, we observed that *XBP1* and *IGHM* showed high gene expression correlation within clone (r = 0,45 and r= 0.41 respectively) (**Figure 3i**), In contrast, when we analysed random B cell pairs from the same donor, there was no significant correlation (**Figure 3i**). This indicated that the level of antibody expression is similar between sister cells. Given that XBP1 is a transcription factor, we reasoned that if its expression is clonally correlated, then the genes it regulates should also show similar correlation patterns. To test this, we examined the expression correlation of XBP1 regulon genes among sister cells.

We found that these genes displayed higher correlation between sister cells than all other genes (**Figure 3j, Supplementary Fig. 3c**), indicating that genes critical to the plasma cell lineage tend to be clonally inherited in their expression profiles. Finally, we asked whether these intra-clonal gene correlations could lead to clone-specific gene expression signatures. To test this, we trained a classifier to predict sister cells based on gene expression patterns and validated it against VDJ-assigned clonal identities. Remarkably, using only 400 correlated genes (**Supplementary Table 5**) we were able to correctly predict sister cells for 80% of the cells (**Figure 3k, Supplementary Fig. 3d**). This indicated that the expression level of a multitude of genes involved in B cell function are highly heritable across divisions, resulting in sister cells with similar gene expression patterns.

Taken together we demonstrated that one naive B cell has the potential to generate both plasma cells and GC cells, and that gene expression levels of genes involved in B cell differentiation are highly clonally regulated.

### Class-switching is clonally independent

Next, we investigated how naive and memory B cells regulate the production of antibodies. As IgM isotype is most common in our culture (i.e. 90% of cells are IgM^+^, **Supplementary Fig. 4a**), specifically, we quantified the ratio of secretory (*sIGHM*) versus membrane (*mIGHM)* transcript isoforms across the time course of activation. As expected, at resting (day 0) and at early activation time point (day 1), both naive and memory B cells predominantly expressed the *mIGHM* (**Figure 4a**). However, by day 3, the isoform ratio had already shifted in favour of the *sIGHM* in memory B cells and not in naive. This trend continued by day 6, when nearly all memory cells expressed *sIGHM* at substantially higher levels than naive cells. In contrast, naive cells retained a more balanced *IGHM* isoform ratio. This difference in isoform ratio dynamics between naive and memory B cells is likely explained by the expression dynamics of *ELL2* (**Figure 4b**), known to promote the use of the secretory-specific polyadenylation site in the *IGHM* gene (Martincic et al. 2009). Notably, *ELL2* expression is regulated by IRF4 (**Supplementary Table 4**), which is exclusively active in plasma cells (**Figure 2b**), consistent with our observation that the secretory isoform was detected in plasma cells but not in GC cells at day 6 (**Figure 4c**).

**Figure 4.**
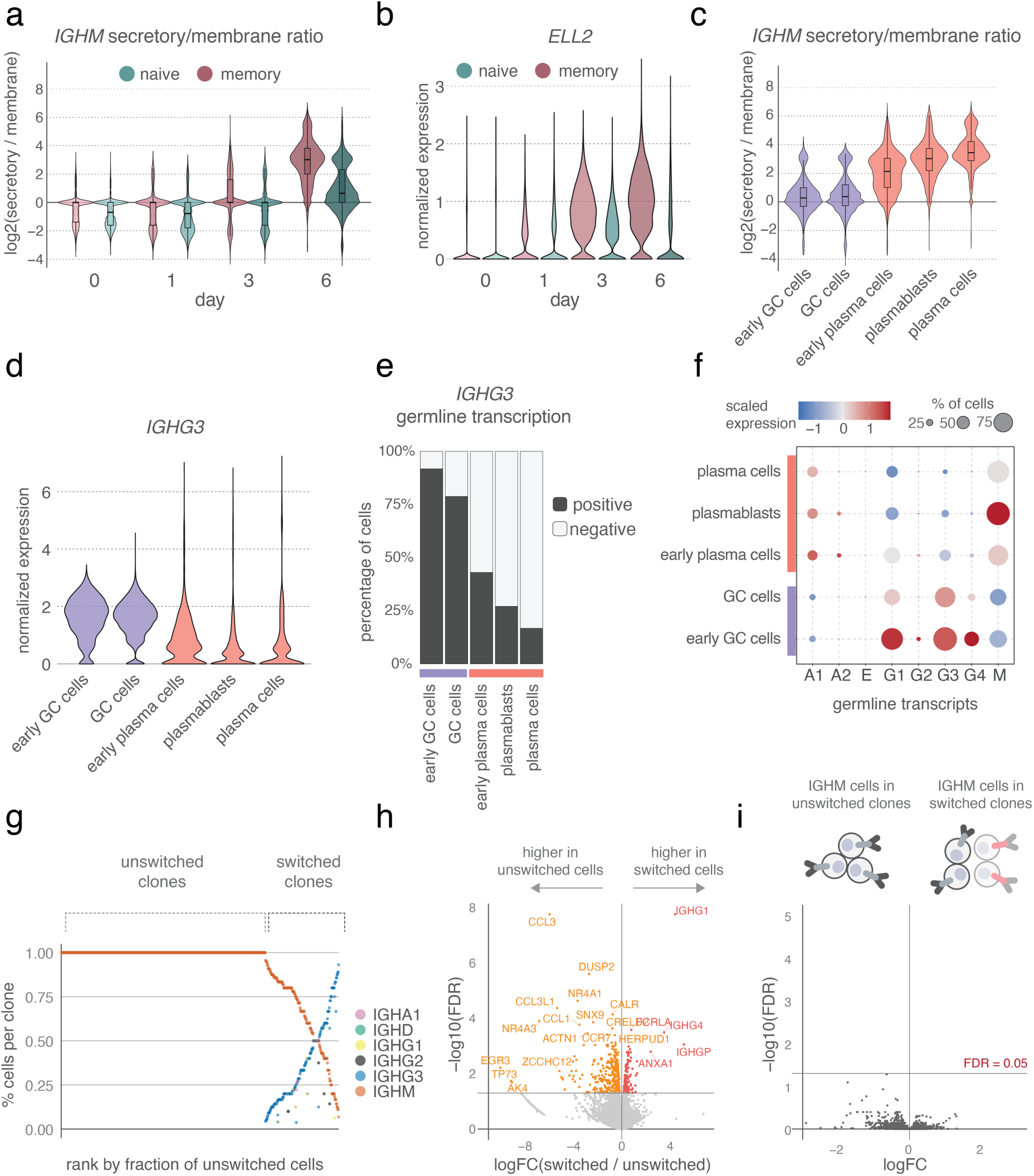
Class-switching is clonally independent. **a**) Ratio of *IGHM* secretory to *IGHM* membrane mRNA isoforms in naive and memory B cells per time point. **b**) Normalized expression level of *ELL2* in naive and memory B cells per time point. **c**) Ratio of *IGHM* secretory to *IGHM* membrane mRNA isoforms in GC and plasma cells. **d**) Normalized expression level of *IGHG3* in GC and plasma cells. **e**) Proportion of cells expressing sterile and productive *IGHG3* transcripts. **f**) Dotplot of GLT expression in GC and plasma cells. Scaled gene expression is represented on a blue-to-red gradient: blue indicates below-average expression, red indicates above-average expression. Dot size reflects the proportion of cells expressing each GLT. **g**) Isotype fractions per clone. Clones are ranked by the fraction of IGHM^+^ cells. Colors represent isotypes. **h**) Volcano plot of differential expression between unswitched and switched cells. Grey represents non-differentially expressed genes, orange and red represent upregulated and downregulated genes in switched cells, respectively. **i**) Volcano plot of differential expression between IGHM^+^ cells from unswitched and switched clones.

Next, we explored isotype switching and found that a large proportion of GC cells expressed *IGHG3* (**Figure 4d**), which was unexpected given that only 15% of day 6 activated naive cells switched antibody class (**Supplementary Fig. 4a**). To investigate this discrepancy, we re-analysed the data by remapping reads using the sciCSR pipeline (Ng et al. 2023), enabling more accurate detection and quantification of antibody isotypes. Isotype switching requires the expression of short transcripts that are transcribed upstream of the transcription start site of the first exon of every isotype, known as germline transcripts (GLTs) (Chaudhuri et al. 2003; Z. Xu et al. 2012; Lorenz, Jung, and Radbruch 1995; Chowdhury et al. 2008). This transcript does not result in a functional antibody protein, but it is required in the recruitment of AID during the class-switching process, thus marking the switching regions poised for CSR. Strikingly, above 75% of GC cells expressed germline *IGHG3* while only 25% of plasma cells showed this pattern (**Figure 4e**). This priming, however, did not lead to an increase in the proportion of switched cells in GC (**Supplementary Fig. 4b**), as GLTs are necessary but not sufficient to lead to antibody class switching. We observed a similar pattern for *IGHG1* and to a lesser extent for *IGHG4* (**Figure 4f**). Therefore, together this demonstrated that germline transcription in the *IGH* antibody locus is significantly more active in GC cells than in plasma cells, suggesting a cell-state specific transcription activity in the IGH locus.

To investigate potential transcriptional differences between switched and unswitched B cells, we focused on characterizing cells that had undergone CSR during *in vitro* stimulation. Although switched cells were excluded during the initial sorting of naive and memory populations, some may still be present due to low-level contamination. To confidently distinguish between contamination and CSR upon activation, we leveraged clonal information and included only clones containing at least one *IGHM*-expressing cell. Since CSR is one-directional and irreversible, the presence of *IGHM* within a clone ensured that any observed isotypes arose from activation rather than from pre-existing switched cells in the starting population. Using this approach, we identified 764 clones containing only unswitched cells and 258 clones containing switched cells in varying proportions (**Figure 4g**). We then performed differential gene expression analysis and detected 196 upregulated and 377 downregulated genes in switched cells (**Figure 4h, Supplementary Table 6**), indicating that CSR is associated with distinct transcriptional programs. Interestingly, we observed higher ER-stress related signatures, including “Protein processing in endoplasmic reticulum” in unswitched cells (**Supplementary Fig.4c, Supplementary Table 7**) which may be explained by the fact that due to the size and complexity of multimeric IgM, its protein folding is more complex than the one of IgG. Finally, a mathematical model of isotype switching demonstrated recently that CSR is independent of clonal membership (Horton et al. 2022), suggesting an underlying stochastic process. This would, in turn, imply that there is no transcriptional determinant of switching, and a probability of switching of a single IgM^+^ cell is independent of its sister cell. To test this further, we investigated the transcriptional difference between IgM^+^ cells from clones containing switched cells, and IgM^+^ cells from unswitched clones. We observed no transcriptional differences between them (**Figure 4i**), demonstrating that IgM^+^ cells from switched clones are not different to IgM^+^ cells from unswitched clones. This implies that belonging to the same clonal ancestry as a switched cell, does not increase likelihood of further switching, validating proposed mathematical model.

### PRDM1 and IRF4 have different roles in fate bifurcation

Since we observed that IRF4 activity has distinct dynamics during naive and memory B cell differentiation trajectory, we next investigated how it influences cell fate decisions and antibody production in both cell types. IRF4 is a well-known regulator of transcription factor PRDM1 driving plasma cell differentiation. To examine their role in both naive and memory B cell differentiation, we knocked out IRF4 and PRDM1 (Methods). This resulted in a marked reduction in the frequency of plasma cells across both cell types, though the effect of IRF4 KO appeared to be stronger (**Figure 5a**). This was accompanied by a significant decrease in the secretory to membrane *IGHM* isoform ratio (**Figure 5b**). As expected the cells that were KO for IRF4 exhibited significant reduction in IRF4 activity, evident by decrease in IRF4 regulon scores (**Figure 5c**). Strikingly, instead they all showed increased activity of the SPI1 (PU1) regulon, which has not been observed in control memory cells (**Figure 5c**). A similar effect was observed following PRDM1 knockout, though the effect on memory was less pronounced than on naive B cells (**Figure 5c**). Finally, we observed a strong increase in germline transcription of IgH genes with the particular increase in germline *IGHG1*, *IGHG1*, *IGHG3* and *IGHA1* (**Figure 5d**).

**Figure 5.**
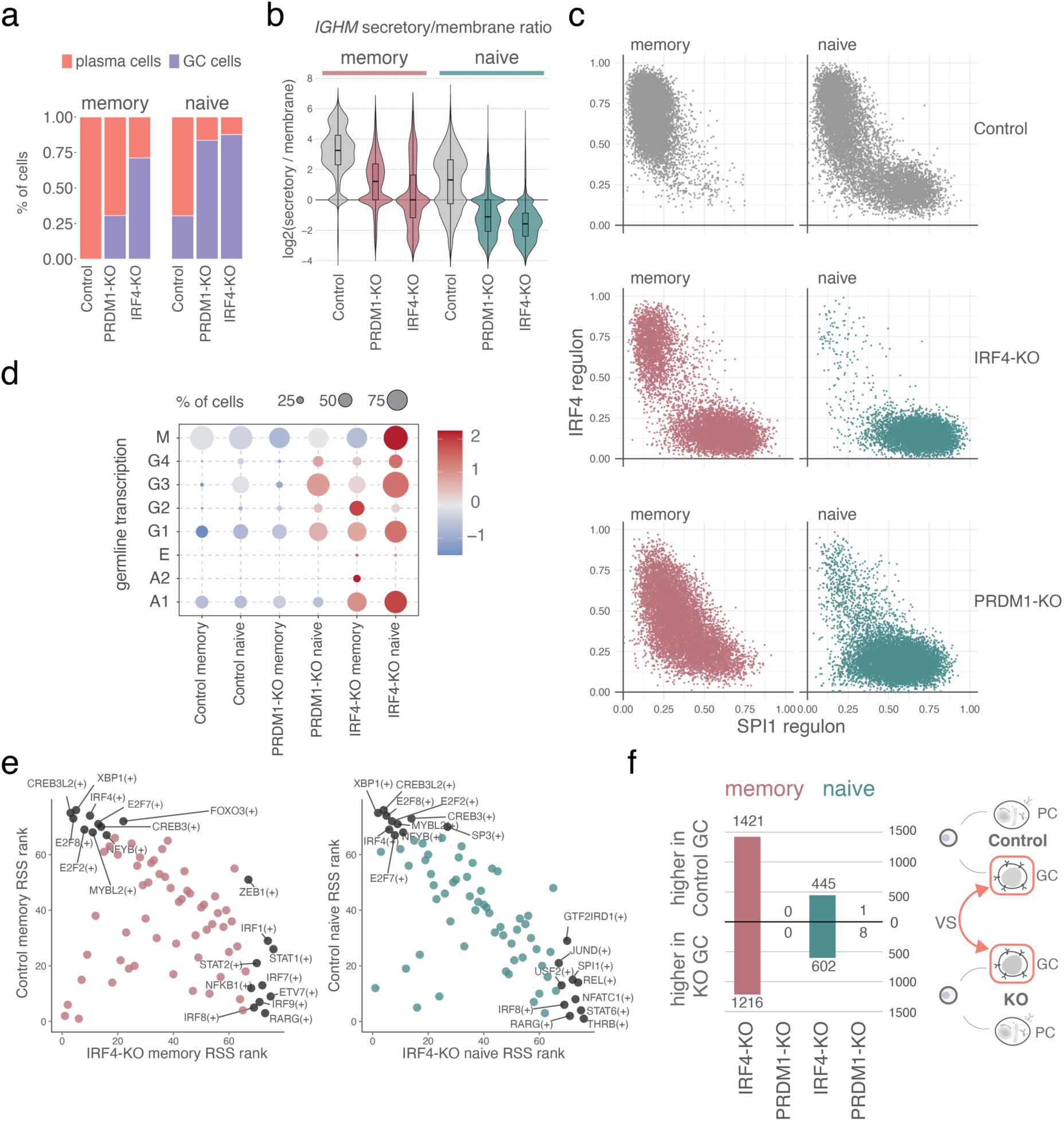
PRDM1 and IRF4 have different roles in fate bifurcation. **a**) Proportion of cells in different cell states for naive and memory cells at day 6. Purple and orange represent GC cells and plasma cells, respectively. **b**) Ratio of *IGHM* secretory to *IGHM* membrane mRNA isoforms in naive and memory B cells per condition. **c**) Dotplot showing the SPI1 and IRF4 regulon scores in naive and memory B cells at day 6. **d**) Dotplot of germline transcription expression in day 6 naive and memory cell knock-out and control conditions. Scaled gene expression is represented on a blue-to-red gradient: blue indicates below-average expression, red indicates above-average expression. Dot size reflects the proportion of cells expressing each GLT. **e**) Comparison of RSS ranks between IRF4-KO and control conditions in naive and memory B cells. For each condition and cell type, RSS are ranked from lowest to highest. Pearson correlation coefficient (r) and its two-sided test P values are shown. Top 10 highly ranked regulons per condition are highlighted in black. **f**) Number of differentially expressed genes between naive control GC-cells and all other conditions.

In addition, we noted substantial transcriptional rewiring following IRF4 and PRDM1 KOs, with over 5000 and 2000 genes differentially expressed compared to control respectively (**Supplementary Table 8**). However, both IRF4 and PRDM1 KOs caused a shift in GRN activity compared to control, as indicated by large deviations in regulon specificity scores from the diagonal (**Figure 5e**). For instance, the XBP1 regulon which is highly active in control condition was completely inactive following IRF4 and PRDM1 KO (**Figure 5e**). Next, we investigated whether the emergence of SPI1-regulon high cells in activated memory B cells following IRF4 and PRDM1 KOs resembled the GC cells arising from natural naive B cell differentiation. To investigate this, we compared GC cells from control naive cells to those arising in memory cells following IRF4 and PRDM1 KOs. Strikingly, GC cells from memory B cells lacking PRDM1 were transcriptionally indistinguishable from those arising from naive cells in control condition (**Figure 5f**). This demonstrated that PRDM1 is only essential for plasma cell generation but dispensable for GC cell development. In contrast, GC cells emerging in memory cells following IRF4 KO displayed over 2637 differentially expressed genes compared to naturally arising GC cells following activation of naive B cells (Supplementary Table 9). Despite expressing canonical GC markers (Supplementary Fig. 5a), these cells were transcriptionally highly distinct, indicating that IRF4 plays a role in both plasma cell and GC differentiation (**Figure 5f**). This suggests that while the plasma cell differentiation was blocked, GC differentiation was impaired, resulting in an aberrant GC state.

Together, our findings support a model (**Figure 6**) in which steady increase of IRF4 activity during activation in memory B cells drives PRDM1 and exclusive differentiation into plasma cells. This activity of IRF4 appears to suppress germline IgH transcript production, thereby reducing the probability of class switching. In naive B cells, however, IRF4 activity increases during early activation but then declines, remaining active only in the subset of cells that commit to the plasma cell fate. Since our system relies on polyclonal stimulation, we can exclude that this is a consequence of cell selection. Thus this bifurcation likely results from stochastic oscillations in IRF4 expression. Finally, as IRF4 has been shown to interact with PU.1 (SPI1) and drive GC fate, its absence may allow PU1 activity to somewhat drive GC identity, however, the resulting cells are not equivalent to GC cells, highlighting the indispensable role of IRF4 in coordinating normal GC and plasma cell differentiation. This role of IRF4 is independent of PRDM1, as evident by the fact that PRDM1 KO does not impair GC cells.

**Figure 6.**
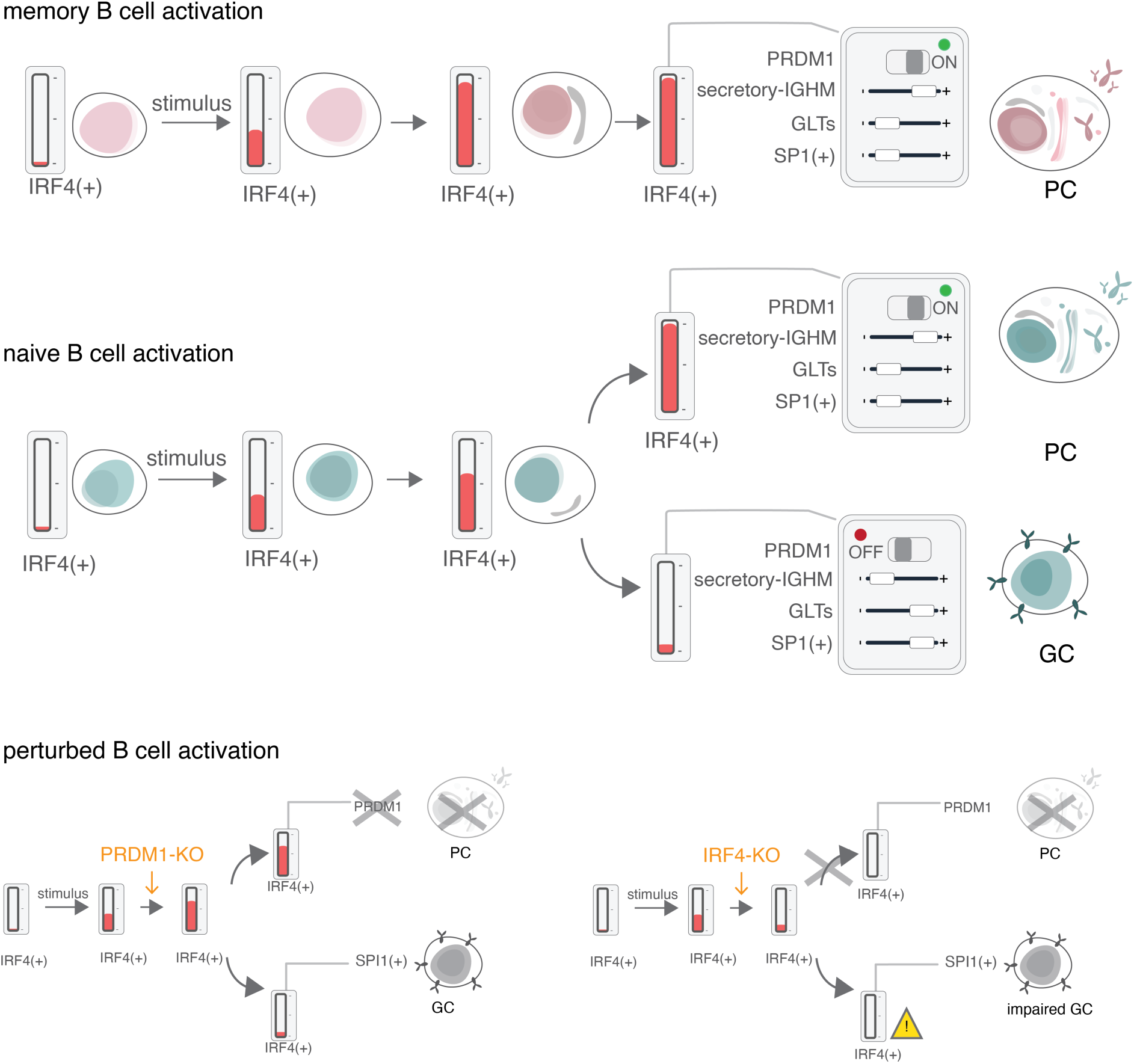
**Model of IRF4 control of naive and memory B cell differentiation.**

## Discussion

Charting the GRNs that control naive and memory B cell differentiation is critical for understanding immune responses to infection, vaccination and autoantigens. In this study, we investigated the cell fate decisions made by human naive and memory B cells. Strikingly, memory B cells almost exclusively differentiated into plasma cells, while naive B cells had a bifurcated differentiation and generated two distinct cell states. One differentiation branch resulted in plasmablasts and plasma cells that clustered with memory counterparts, while the other resulted in a germinal centre (GC) phenotype. By analyzing gene expression dynamics over six days of stimulation, we mapped the GRNs active at different stages of B cell activation. We showed that memory B cells had a more robust IRF4 activity upon activation than naive B cells, a phenomenon already observed in the first 24h of activation. Previous work in mice demonstrated that transient IRF4 expression results in co-binding with PU.1 (SPI1) at AICE motifs to promote GC formation, but sustained expression results in shifts to lower-affinity ISRE motifs associated with plasma cell genes (Ochiai et al. 2013; Shao et al. 2024). In line with this we demonstrated that IRF4 is a key network responsible for more efficient differentiation of memory than naive B cells towards plasma cells. On the other hand, in naive B cells, its expression rises more slowly during early activation but then declines, persisting only in those committing to the plasma cell fate. Because our polyclonal stimulation system excludes selection effects, this bifurcation likely arises from stochastic fluctuations in IRF4 expression.

We hypothesized that knocking out PRDM1 and its upstream regulator IRF4 would rewire the B cell transcriptome, blocking plasma cell differentiation and favoring a GC fate. Interestingly, in IRF4-knockout cells, both naive and memory B cells gave rise to a population resembling GC cells, although this state was notably distinct from the GC cells observed in controls. We therefore hypothesise that this state represents an aberrant GC differentiation. The fact that IRF4 is essential not only for plasma cell differentiation but also for aspects of the GC program aligns with previous observations that IRF4 can dimerize with PU.1 (SPI1) to activate BACH2 transcription and induction of GC genes (Carotta et al. 2014; Ochiai et al. 2024). Consistently, we observed an anticorrelated activity pattern between the IRF4 and SPI1 (PU.1) regulatory networks, with SPI1 (PU.1) activity completely silenced during the terminal differentiation of naive B cells into plasma cells. On the other hand, PRDM1-knockout naive and memory B cells had impaired plasma cell generation as expected, but strikingly produced GC cells highly similar to those in the control condition. These results demonstrate that although memory B cells are intrinsically biased toward plasma cell differentiation, they retain the capacity to adopt a GC fate when PRDM1 activity is suppressed, opening new avenues to direct memory cells to GC state.

Stochastic fluctuations in the expression of transcription factors at the single-cell level could explain why in our clonal tracking experiment, cells originating from the same ancestral cell (i.e. a clone) were observed to adopt both PC and GC fate. This finding aligns with previous observation in mice, where one cell can give multiple fates (Taylor et al. 2015). The authors showed that the ability of B cells to differentiate into these subtypes was linked to the affinity of their BCRs. In our system, we demonstrated that even in the absence of antigen affinity difference, one naive B cell has the plasticity to take both roads of differentiation. Interestingly, we found a correlation between clonal size and clonal composition - smaller clones tended to have a higher proportion of GC cells, suggesting that proliferation capacity may vary even among cells from the same clone.

Although naive B cells can divide and differentiate into multiple fates, cells within the same clone exhibited sharing in their gene expression profiles. In fact, gene expression alone was sufficient to identify clonal families. This indicates that, even under polyclonal stimulation and in the absence of selection, individual clones diverge from one another. Heritable gene expression patterns within clonal families have also been reported in other immune cell types, such as T cells and NK cells (Rückert et al. 2022; Mold et al. 2024). These findings support the hypothesis that the final composition of cell subsets within a clone may be an intrinsic property of that clone. One can further hypothesize that initial differences in signalling dynamics during activation, oscillation in IRF4 expression, or proliferation kinetics may contribute to shaping the ultimate distribution of B cell fates within each clone, adding an additional layer of variability to the B cell immune response.

Interestingly, when comparing cell populations, over 75% of GC cells expressed germline transcripts, while this was true for only about 25% of plasma cells. Despite this strong difference in GLT expression, there was no corresponding difference in the proportion of cells that had undergone class switching between the two populations. This aligns with the fact that while GLT expression is necessary for class switching, it is not sufficient to drive the process. Moreover, we observed that *IRF4* knockout cells had a marked increase in GLT levels. This implies that high IRF4 activity may act to suppress GLT expression, adding a potential regulatory role for IRF4 in controlling the accessibility of IgH regions.

Finally, this led us to investigate whether isotype switching is clonally regulated by comparing unswitched cells from both switched and unswitched clones. We found no transcriptional differences between the two groups. Ultimately, this implies that being part of a clonal lineage that has already undergone switching does not confer an increased predisposition toward further switching. These results support the idea that isotype switching is a stochastic and cell-intrinsic process, rather than one governed by a shared transcriptional trajectory within a clone. This independence for CSR among clonally related cells aligns with and provides further experimental evidence for models (Horton et al. 2022) that describe isotype switching as a probabilistic process occurring independently across individual cells within a clone. Our results also likely explain why isotype switching in early GC B cells is highly variable *in vivo*.

Taken together, our findings support a model in which each individual cell is governed by a unique combination of gene programs that regulate cell state, clonal gene expression, and isotype switching, maximizing the breadth of diversity in the immune response.

## Methods

### Naive and memory B cell isolation

Human biological samples were sourced ethically, and their research use was in accordance with the terms of the informed consents under an approved protocol (ID 3441, study number 6619). Peripheral blood mononuclear cells (PBMCs) were isolated from healthy blood donors using Ficoll-Hypaque density gradient centrifugation. Prior to B cell isolation, PBMCs were incubated overnight (12 hours) at 5 million cells/mL in RPMI 1640, HEPES and glutamine supplemented with 10% fetal bovine serum and penicillin/streptomycin (ThermoFisher Scientific). Total B cells were then isolated by negative selection using the EasySep™ Human B Cell Isolation Kit (Stemcell Technologies). B cells were stained in DPBS with eFluor780 Live/Dead dye (Thermo Fisher Scientific) for 10 minutes at 4°C according to manufacturer’s instruction followed by washing with FACS buffer (1 x DPBS, 3% FBS, 1mM EDTA (Merck)) and staining with surface fluorochrome-labeled primary antibodies for 20 minutes at 4°C. The following antibodies were used: anti-human CD19-BV786 (BD Bioscience), anti-human CD27-VioBright B515, CD20-APC, IgG-PE, and IgA-PerCP-Vio700 (Milteny). Cells were washed with HBSS, 2% FBS, 10 mM Hepes resuspended in the same buffer and sorted using a MoFlo Astrios EQ cell sorter (Beckman Coulter, USA).

### In vitro B cell activation

Sorted naive and memory B cells were cultured in complete StemPro-34 SFM medium supplemented with 2mM glutamine and 100 U of Pen/Strep. 25.000 cells were seeded in 600 μl of media in 48 well plate wells. Sorted B cells were activated in complete StemPro-34 medium with 200 ng/mL megaCD40L (Enzo Life Science), 5µg/mL AffiniPure F(ab’)2 Fragment Goat Anti-Human IgM, Fc5 Fragment Specific (Jackson ImmunoResearch), 100 ng/mL IL-21, and 20 ng/mL IL-2 (PeproTech, ThermoFisher Scientific) at 37°C. At the designated time points, live cells were sorted in for the scRNA-seq. For scRNAseq we independently activated naive and memory B cells from four healthy donors.

### Clonal culture

Naive B cells were isolated and sorted as described above. 2000 naive B cells (live, CD19⁺ CD20⁺ CD27⁻ IgG⁻ IgA⁻) from two donors were sorted directly into 96 round-bottom well plates. Cells were cultured in 200μl in complete StemPro-34 medium, and B cells were stimulated as described above. At day 6, cell count and viability were determined with the Trypan blue staining.

### CRISPR-CAS9 gene knock-outs

Synthego’s *IRF4, PRDM1*, and *AAVS1* (within *PPP1R12C* gene) Gene Knockout Kits v2, along with Cas9 2NLS, were supplied by EditCo Bio. Briefly, naive and memory B cells were sorted from pre-enriched total B cells of two healthy volunteers, as previously described, using a MoFlo Astrios cell sorter (Beckman Coulter). Sorted B cells were activated in complete StemPro-34 medium with 200 ng/mL megaCD40L (Enzo Life Science), 5µg/mL AffiniPure F(ab’)2 Fragment Goat Anti-Human IgM, Fc5 Fragment Specific (Jackson ImmunoResearch), 100 ng/mL IL-21, and 20 ng/mL IL-2 (PeproTech, ThermoFisher Scientific) at 37°C for 20 hours. Cells were collected and electroporated at 1 x 10 ^5 cells for each target gene with the sgRNA-Cas9 mixture at a 9:1 ratio using the P3 Primary Cell 4D-Nucleofector X Kit (Lonza) following the manufacturer’s instructions and rested in complete StemPro-34 medium at 37°C for 30 minutes. Naive and memory B cells were activated as described above and cultured for a further 5 days. On day 6, cells were collected and processed for scRNA and paired with scBCR using the 10X Genomics Chromium Next GEM Single Cell 5’ v2 Kit. The efficiency of CRISPR was assessed by flow cytometry and qualitative PCR. sgRNA multi-guide sequences, provided by Editco. Bio, are: huIRF4: CACGCGGGGCAUGAACCUGG, GCGCGGUGAGCUGCGGCAAC, AGAGCAUCUUCCGCAUCCCC; huPRDM1: GUUGGCAGGGAUGGGCUUAA, GAAGUGGUGAAGCUCCCCUC, CUCUCCCCGGGAGCAAAACC; PPP1R12C: CUCCAGGUUCUCAUCAAUGC, GUGGCUACCUAGAUAUCGCC, GUUGUCUGCCUGGUUCACAG.

### PCR analysis of Gene knockout

DNA was isolated from human B cell pellets using QIAamp UCP DNA mini spin columns (Qiagen), following the manufacturer’s instructions. EditCo.Bio-supplied PCR primer sequences were utilised for the PCR: huIRF4: TCGTGGTCACTGGCGCA, ACGCCACCTGATGCCTC, amplicon size 500 bp; huPRDM1: CGCCCTGATTTCTGCTGATTC, CATGTTATTAGTTCAAAGGGGCAG, amplicon size 500 bp; PPP1R12C:

CTTCAGCAGCCCCTCCATG, CCAGGCGTATCTTAAACAGCC, amplicon size 475 bp. Briefly, 100 ng of isolated DNA was amplified using the Platinum SuperFi PCR ready-to-use master mix (Thermofisher Scientific) according to the manufacturer’s instructions. Samples were processed using the VerityPro 96-well thermocycler from Applied Biosystems, using the following cycling conditions: 98 °C for 30 s, 30 cycles of 98 °C for 10 s, 60 °C for 10 s, 72 °C for 15 s, and 72 °C for 5 min. Amplicons were visualised on a 2% agarose gel.

### Flow cytometry

For surface staining, cells were washed with PBS and incubated with Live/Dead Fixable Near IR (876) Viability dye (ThermoFisher Scientific), diluted 1:2000 in PBS, for 15 minutes on ice. Cells were then washed with a staining buffer (PBS, 3% FBS, and 0.05% Sodium Azide) before being incubated for 30 minutes at 4 °C with an antibody mix targeting cell surface markers. Cells were then fixed with IC Fixation Buffer (ThermoFisher Scientific) for 1h at 4°C. For intracellular markers, cells were permeabilized using either the Transcription Factor Buffer Set (BD Bioscience) or the True-Nuclear Transcription Factor Buffer Set, in accordance with the manufacturer’s instructions (Biolegend). Cell Trace Violet was prepared according to the manufacturer’s instructions (Thermofisher Scientific), and the solutions were warmed to 37°C. An equal volume of CTV in PBS was added to the cell suspension to achieve a final concentration of 2.5 µM dye (1:2000). The incubation was performed at 37°C and 5% CO2 for 20 minutes, protected from light. The dye was quenched by adding three times the original volume of RPMI 1640 HEPES-Glutamine medium, which contained 10% FBS, 100 U of penicillin/streptomycin, followed by a 5-minute incubation at RT. Samples were run on a CytoFLEX LX (Beckman Coulter). Data analysis was performed using the FlowJo data analysis software package (TreeStar, USA). Immunophenotyping included: ZombieNir (849/876nm) live-dead, CD19-BV786, CD27 Vio Bright B515, IgD Vio Bright R720, CD20-APC, IgG-PE, IgA-PerCPVio700, IgM-BUV395. Data were analyzed with FlowJo and were imported in R (v4.3.1) for statistical analysis.

### Intracellular staining and ImageStream Analysis

Naive B cells were isolated by negative selection using magnetic beads (StemCell Technologies) from PBMC after overnight resting in complete RPMI 1640, HEPES medium (Thermofisher Scientific) supplemented with 10% FBS and 100 µg/mL of penicillin and streptomycin. 1 × 10^5^ cells per well in a 24-well plate were activated with the previously described B cell stimulation cocktail for 6 days in complete STEMPRO-34 medium. On day 6, 5 × 10^5^ cells were collected, washed in PBS, and stained with Fixable Viability Dye eFluor 780 (ThermoFisher Scientific) for 10 minutes at 4 °C. The cells were washed with staining buffer and surface stained for 30 minutes with anti-human CD20 Viobright B515 (Miltenyi), followed by intracellular staining with anti-human/mouse IRF4 eFluor 450 (ThermoFisher Scientific) and anti-human PU.1 (SPI1) PE (Biolegend) monoclonal antibodies using the True-Nuclear Transcription Factor Buffer set (Biolegend) as per the manufacturer’s instructions. Briefly, the cells were fixed in True-Nuclear 1x-Fix Buffer at room temperature for 45 minutes, washed twice with True-Nuclear 1x-Perm Buffer and then resuspended in 100 µl of 1x Perm Buffer. After blocking for 5 minutes with Human TrueStain FcX (Biolegend), the conjugated antibodies were added and the sample was incubated for 30 minutes at room temperature. At the end of the incubation, 0.25 µl of DRAQ5 nuclear staining (Thermofisher Scientific) in 500 µL of 1x Perm was added to the sample for 5 minutes. The sample was washed with 1X perm and resuspended in 25 µL of Staining Buffer. Sample acquisition was performed using the Cytek Amnis ImageStream MkII Flow Cytometer; data analysis was conducted with the IDEAS Image Analysis Software.

### scRNA-seq library preparation

At each time point, naive and memory B cells were collected and were stained with eFluor780-Live/Dead. Live cells were sorted with the MoFlo Astrios FACS sorter. Naive B cells and memory B cells deriving from two different donors and from the same time point, were pooled together in one single cell 10x reaction. This single cell experiment configuration was replicated for each time point of the analysis. Single cells were encapsulated into Gel Emulsion droplets using the Chromium Controller system (10x Genomics) targeting 20’000 cells for each sample and 5’ Gene Expression and BCR libraries were generated in accordance with the protocol ChromiumNextGEMSingleCell5-v2 CG000331 (10x Genomics). Libraries were pooled at a ratio dependent on the 10x Genomics indication and loaded on the Novaseq6000 sequencer to achieve a minimum of 50’000 reads/cell for each Gene Expression library and 5000 reads/cell per BCR library.

### Genotyping for donor deconvolution

DNA was extracted from each donors’ PBMCs. Low-Pass whole genome sequencing (0.5-1X) was performed with Illumina NovaSeq600. Genotypes were inferred with the sarek nf-core pipeline (Ewels et al. 2020; ‘Sarek: Introduction’, n.d.). Genotype imputation was performed with GLIMPSE2 (Rubinacci et al. 2021).

### scRNA-seq preprocessing and clustering

Sequencing reads were aligned to the hg38 reference genome and quantified with CellRanger (v7.0.1). Individual samples were demultiplexed by first genotyping each cell with cellSNP-lite (X. Huang and Huang 2021), and genotype deconvolution was performed with VireoSNP by providing donor genotypes (Y. Huang, McCarthy, and Stegle 2019). Background RNA was removed from the count matrices with cellbender (v0.2.0)(Fleming et al. 2023). Quality control, doublet removal, filtering, clustering, dimensionality reduction, data inspection was performed in python (v3.12) using the Scanpy package (v1.10.1). Quality control was performed filtering out cells with low number of counts, low number of genes and high fraction of mitochondrial genes using adaptive thresholds (‘Orchestrating Single-Cell Analysis with Bioconductor’, n.d.; Heumos et al. 2023). Doublets were identified and removed with scrublet (‘Scrublet: Computational Identification of Cell Doublets in Single-Cell Transcriptomic Data’ 2019). Counts were first normalized over the total counts over all genes, then multiplied by 10000 and lastly the data was log transformed. Before dimensionality reduction, we removed from the variable features all immunoglobulin variable genes as well as the heavy constant and light constant chains. Dimensionality reduction was performed with principal component analyses and cells were clustered based on the first 30 principal components. Quality control was repeated after the first clustering iteration as a cluster of likely dying cells was identified and removed before re-performing PCA and clustering. Several clustering resolutions were inspected and we selected the lowest clustering resolution that resolved naive and memory B cell types at each time point. Donor-dependent effects were integrated using Harmony (Korsunsky et al. 2019). Finally, after quality control, our time course dataset consisted of 118.891 cells from four different donors.

### Differential gene expression and pathway enrichment

Differential gene expression was performed using the edgeR package (v4.10.16). Anndata objects were converted to Seurat (5.0.1) objects in R (4.3.1). UMI counts were pseudo-bulked per donor and condition and genes were tested if their aggregated sum was at least 30 UMIs. Differential expression was tested with the glmTreat method by testing if the genes were differentially expressed by at least 20% between conditions accounting for the donor effect. To identify relevant pathways enriched in the different conditions tested we performed overrepresentation analysis using the goprofiler2 (v0.2.3) R package, and tested for enrichment in pathways from the GO, KEGG and Reactome databases. Competitive gene set tests were performed with the camera method in the edgeR package (D. Wu and Smyth 2012) using GO, KEGG and Reactome databases.

### scVDJ-seq preprocessing and clonal analyses

scVDJ libraries were aligned with CellRanger (v7.0.1). The Immcantation and (Gupta et al. 2015) and the Dandelion (Suo et al. 2023) frameworks were used for the identification of the alleles, isotypes and to assign cells to clonal families. We started from the unfiltered contigs generated from cellranger and we used the dandelion to run IgBlast to genotype the VDJ alleles in each cell. When cells with multiple VDJ contigs were found, for heavy and light chains, we retained the contig with the highest expression if its expression was three times higher than the summed expression of the other contigs in the same cells. Otherwise the cell was marked as “no contig” and was not considered for further VDJ and clonal analysis. Additionally, chains with less than 3 UMIs were filtered out. Cells were grouped into clonal families using the Immcantation pipeline. We measured Hamming distance between all VDJ-junctions and calculated a threshold for grouping VDJ-sequences into the same clonal family. Thresholds were calculated per donor, and the mean of the threshold were used as the two donors yielded highly similar thresholds (0.12, 0.08). Cells were then grouped into clonal families if they shared the same VDJ alleles. If two cells had the same heavy chain but different light chain, the cells were further separated into two different clonal families.

For the gene expression analysis of the clonal dataset preprocessing and quality control was performed as before. Finally, after quality control, our clonal dataset consisted of 23,484 naive B cells from two different donors. For intraclonal and interclonal correlation, only clones with at least 4 cells were considered and at least one cell had to be *IGHM+*. Interclonal and intraclonal correlation were calculated separately per donor. Only genes expressed in at least 300 cells were used. V(D)J genes were excluded. 10.000 random pairs of cells were randomly sampled from clones and the Pearson correlation coefficient was calculated for each interclonal and intraclonal pairs. To train the classifier we took clones with at least 25 cells. Each clone was divided randomly in 70% cells used for training and 30% used for the validation. A feed-forward neural network was used, with a total of 4 hidden layers. The rectifier linear unit was used as the activation function and dropout of 0.4 was used to avoid overfitting and improve generalization. For the input layer, we first selected the genes whose interclonal correlation was above r of 0.25 and expressed in at least 40% of cells for each donor. The union of the top correlated genes of the donors was then used. Crossentropy was used as a function to calculate the loss. 200 epochs of training were performed on the training dataset. As control, clonal assignments were randomized per donor and the training and validation procedure was repeated as before.

### Isoform quantification with salmon alevin

For transcript-level quantification we used Salmon/Alevin (v1.10.2) (Srivastava et al. 2019). And for differential transcript usage we used DTUrtle (v1.0.2) (Tekath and Dugas 2021).

### Gene regulatory network inference

GRNs were inferred with SCENIC (Aibar et al. 2017; Van de Sande et al. 2020). To reduce the computational burden and reduce the dropout due to sparsity of scRNA-seq data, we performed GRN inference on metacells. Metacells were generated per each sample independently with SEAcells (Persad et al. 2023). We performed 30 runs of the GRN inference and we retained the regulons that were detected in 80% of the runs. Next, for each regulon, we retained the genes that were assigned to that regulon in 70% of the runs. Regulons with less than 10 genes were discarded. Regulon scores were quantified for each cell with AUCell, which normalizes the output of the regulon scores (Aibar et al. 2017). GRN inference was performed with the docker container provided by the scenic authors using the arboreto implementation (Van de Sande et al. 2020). We refer to regulons as sets of genes and the transcription factor that controls their expression. Regulon scores are obtained for each cell with AUCell. To allow comparison of different regulons we used Regulon Specificity Scores (RSS).

## Acknowledgements

We thank the Human Technopole facilities for their contribution to data generation and processing. Particularly we thank the National Facility (NF) for Genomics and NF for Light Imaging. We thank the infrastructural unit for Flow cytometry. PD and PM are PhD students within the European School of Molecular Medicine (SEMM). This research is supported by institutional funds from Human Technopole.

## Author contributions

Conceived and designed the project: PD, BS; Experimental work: PD, LE, PM, ER, PF, CP, SB; Computational analysis: PD, DB, EG, BS; Writing – original draft: PD, BS; Writing – review and editing: all authors; Figure design: PD, BS; Supervision: BS; Funding acquisition: BS.

