## Supplemental figures for "Competing gene regulatory networks drive naive and memory B cell differentiation"

### Supplementary Figures

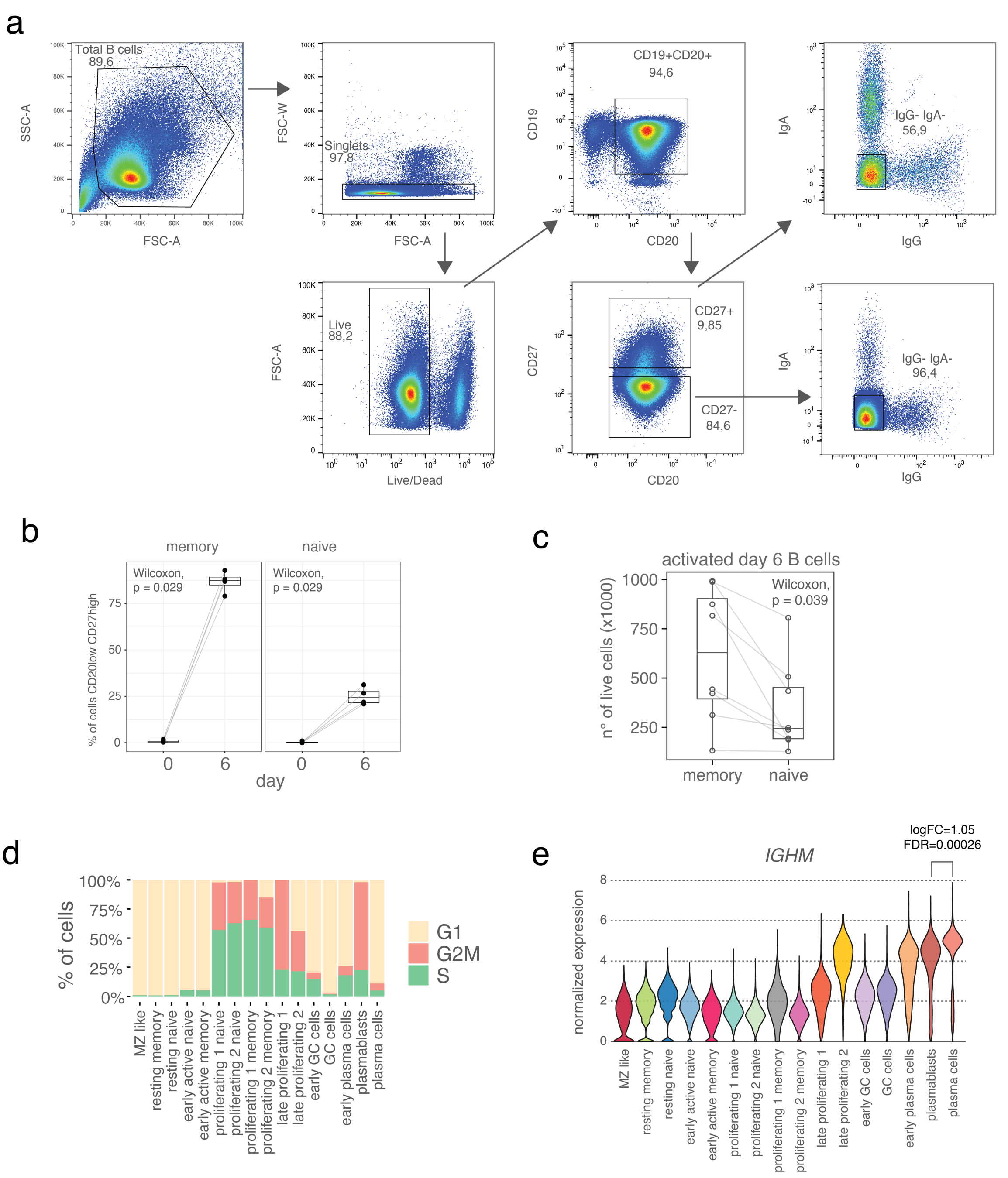


**Supplementary Figure 1 a**) gating strategy for sorting naive (live, CD19$^{+}$ CD20$^{+}$ CD27$_{+}$ IgG$^{-}$ IgA$^{-}$) and memory B cells (live, CD19$^{+}$ CD20$^{+}$ CD27$_{+}$ IgG$^{-}$ IgA$^{-}$) from total B cells. **b**) Fraction CD27$^{high}$ CD20$^{low}$ cells; n=4 biologically independent replicates. **c**) Total number of live cells at day 6; n=8 biologically independent replicates. **d**) Proportion of cells in different cell cycle phases stratified by cell type and time point. **e**) Normalized expression level of *IGHM.* Colors represent cell state.

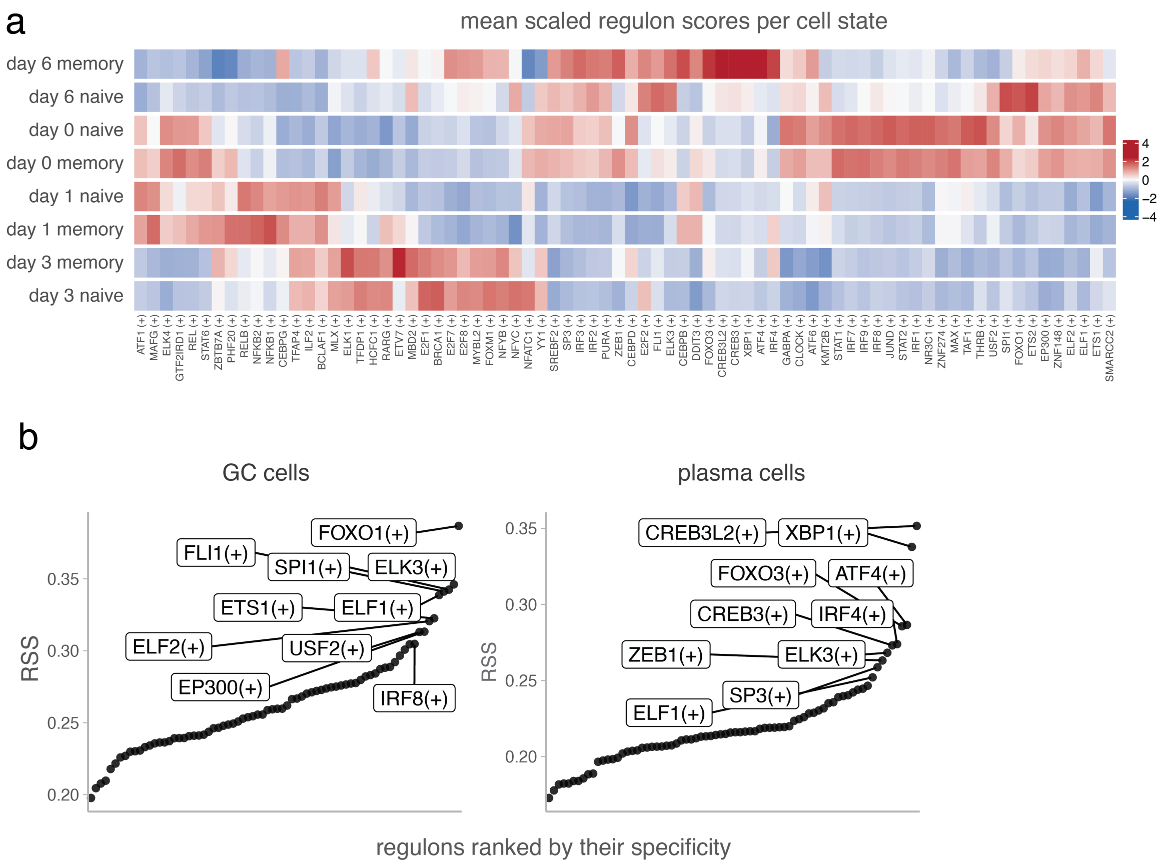


**Supplementary Figure 2 a**) Heatmap of regulon activity per time point and cell type. Columns represent transcription factors. For each condition, mean regulon scores were calculated across all cells and scaled. Blue and red represent score values from lowest to highest. **b**) Regulon specificity scores (RSS) for GC cells and plasma cells. Regulons were ranked by their RSS value in increasing order. For the top 10 regulons, the transcription factor is shown.

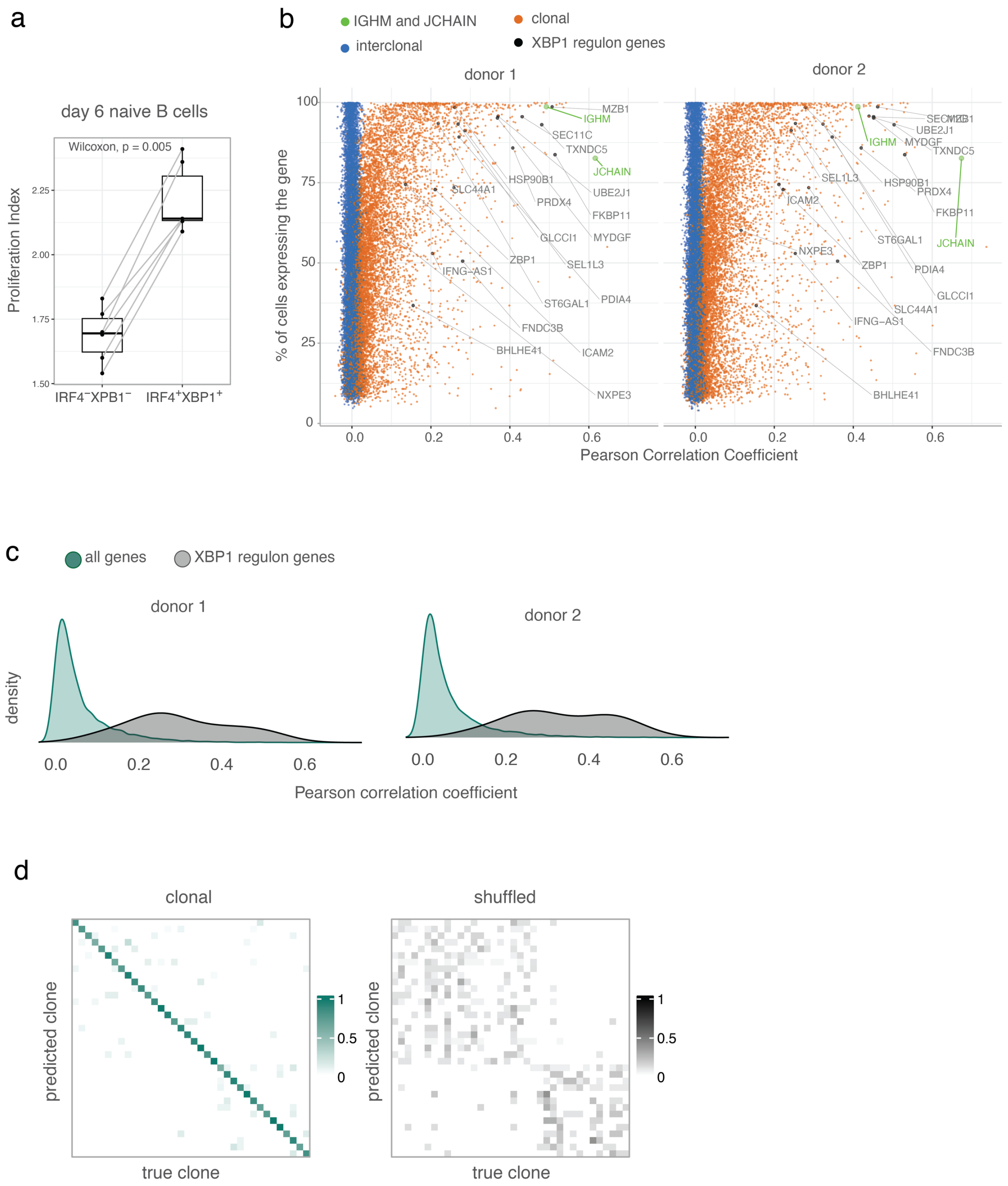


**Supplementary Figure 3 a**) Proliferation index of IRF4⁻XPB1⁻ IRF4⁺XBP1⁺ naive B cells at day 6 post activation, n=6 biologically independent replicates. **b**) Pearson Correlation Coefficient of gene expression against percentage of cells expressing the gene. Each dot represents a gene. Orange and blue represent clonal and interclonal correlations. Genes belonging to the XBP1 regulon are highlighted in black. *IGHM* and *JCHAIN* are highlighted in green. n=2 biologically independent replicates. **c**) Density plot of clonal Pearson correlation coefficient for XBP1 regulon genes compared to all other genes. n=2. **d**) Confusion matrix illustrating the performance of the classifier. Rows represent the predicted clones and columns represent the true clonal assignment. The diagonal shows the proportion of correctly classified clones, while the off-diagonals indicate misclassification rate. Color intensity indicates the frequencies, with darker shades representing higher values. Green bars represent classification using real clonal assignments; grey bars represent a control where clonal groups were shuffled prior to training and validation.

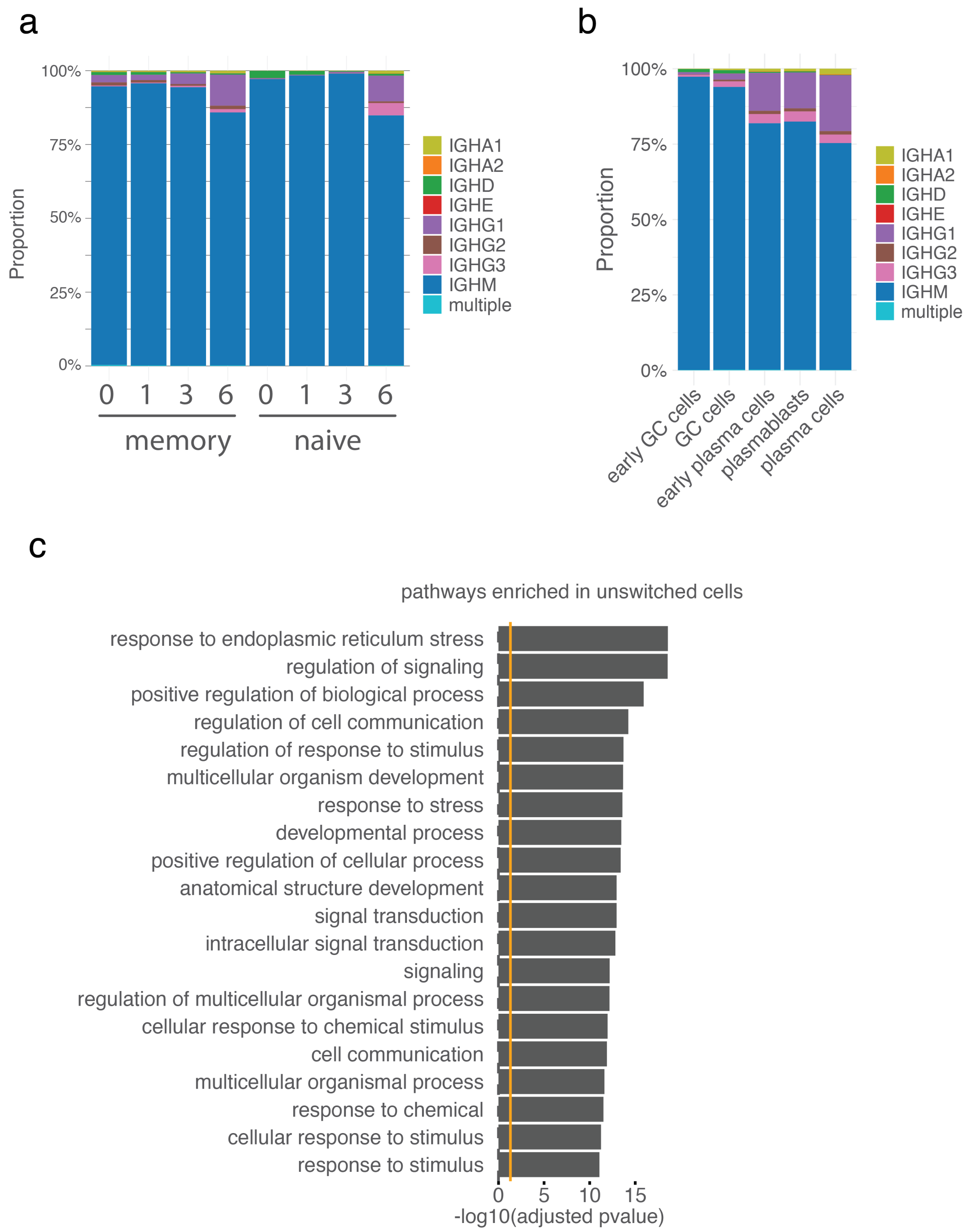


**Supplementary Figure 4 a**) Proportion of antibody isotypes within each time point and cell type group. **b**) Proportion of antibody isotypes within each cell state. **c**) Top enriched reactome pathways in unswitched day 6 clonal naive B cells.

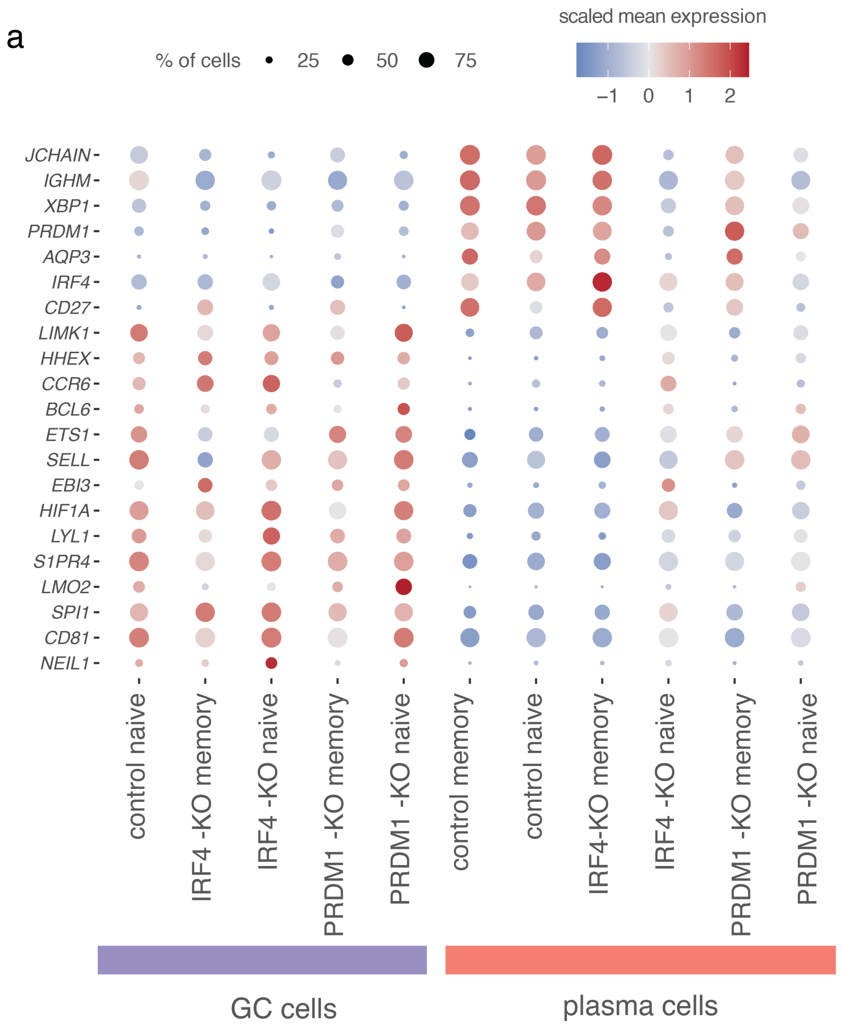


**Supplementary Figure 5 a**) Dot plot of cell state specific genes. Scaled gene expression is represented on a blue-to-red gradient: blue indicates below-average expression, red indicates above-average expression. Dot size reflects the proportion of cells expressing each gene.
